# Efficient prediction of a spatial transcriptomics profile better characterizes breast cancer tissue sections without costly experimentation

**DOI:** 10.1101/2021.04.22.440763

**Authors:** Taku Monjo, Masaru Koido, Satoi Nagasawa, Yutaka Suzuki, Yoichiro Kamatani

## Abstract

Spatial transcriptomics is an emerging technology requiring costly reagents and considerable skills, limiting the identification of transcriptional markers related to histology. Here, we show that predicted spatial gene-expressions in unmeasured regions and tissues can enhance biologists’ histological interpretations. We developed the <u>Deep</u> learning model for <u>Spa</u>tial gene <u>C</u>lusters and <u>E</u>xpression, DeepSpaCE and confirmed its performance using the spatial-transcriptome profiles and immunohistochemistry images of consecutive human breast cancer tissue sections. For example, the predicted expression patterns of *SPARC*, an invasion marker, highlighted a small tumor-invasion region that is difficult to identify using raw data of spatial transcriptome alone because of a lack of measurements. We further developed semi-supervised DeepSpaCE using unlabeled histology images and increased the imputation accuracy of consecutive sections, enhancing applicability for a small sample size. Our method enables users to derive hidden histological characters via spatial transcriptome and gene annotations, leading to accelerated biological discoveries without additional experiments.

## Introduction

Spatial transcriptomics with *in situ* capturing is an emerging technology that maps gene-expression profiles with corresponding spatial information in a tissue section^1–4^. A highly resolved spatial-transcriptome profile is an invaluable resource for revealing biological functions and molecular mechanisms^5^. Recently, many histological transcriptome profiles, measured by *in situ* capturing platforms (numerous spots with barcoded oligonucleotides on a chip), were reported in the field of oncology^6,7^. These profiles have helped demonstrate the complexity and heterogeneity of cancer tissues. For example, histological transcriptome profiles were used to identify high-risk invasive populations in ductal carcinoma tissues using an *in situ* capturing method^8,9^. However, the experimental cost of spatial transcriptomics, such as for designed chips, reagents, and sequencing, is currently high. It is also challenging to maintain the balance between the spatial resolution (i.e., density of spots in a tissue slide) and RNA-detection efficiency with current spatial-transcriptome technology^10^. In addition, this technique requires practiced skills to obtain high-quality expression profiles for entire tissue slides, even when using a commercial kit such as the 10x Genomics Visium platform.

The convolutional neural network (CNN), a deep-learning method, is frequently used for discovering features from imaging datasets and can be used to predict image categories of interest in an end-to-end manner. For example, in the biomedical field, the CNN method has successfully been used to classify lung cancer subtypes from tissue-section images without prior knowledge^11^. Based on these recent advances, we hypothesized that applying the CNN method to spatial-transcriptome profiles would enable expression-level predictions from hematoxylin and eosin (H&E)-stained section images, potentially leading to an increased number of pixels by predicting spatial gene-expression gaps among spots measured by spatial-transcriptome techniques (super-resolution which was inspired by the recent super-resolution technique^12^) or imputing spatial-transcriptomic patterns in unmeasured consecutive sections (tissue section imputation).

Here, we developed the <u>Deep</u> learning model for <u>Spa</u>tial gene <u>C</u>lusters and <u>E</u>xpression (DeepSpaCE), which predicts spatial-transcriptome profiles from H&E-stained images using CNNs. We verified the prediction accuracy of this model by comparing expression profiles from testing datasets and protein-expression patterns in adjacent sections (using immunohistochemistry data), which were consistent with the predictions. Based on these verifications, we applied DeepSpaCE for super-resolution of spatial gene-expression levels and imputation of spatial gene-expression levels in other tissue sections using human breast cancer datasets.

## Results

### Overview of DeepSpaCE

DeepSpaCE is composed of two parts: the training part and gene-prediction part (**Fig. 1**). The CNN (VGG16 architecture) is trained with pairs of cropped section images for each spot (spot image) and its gene-expression profiles. Next, the trained model predicts gene-expression levels for at least one transcript (or transcriptomic cluster type) from spot images. We conducted two types of practical applications of DeepSpaCE using the *in situ* capturing spatial transcriptome dataset: (a) super-resolution and (b) tissue section imputation.

**Figure 1.**
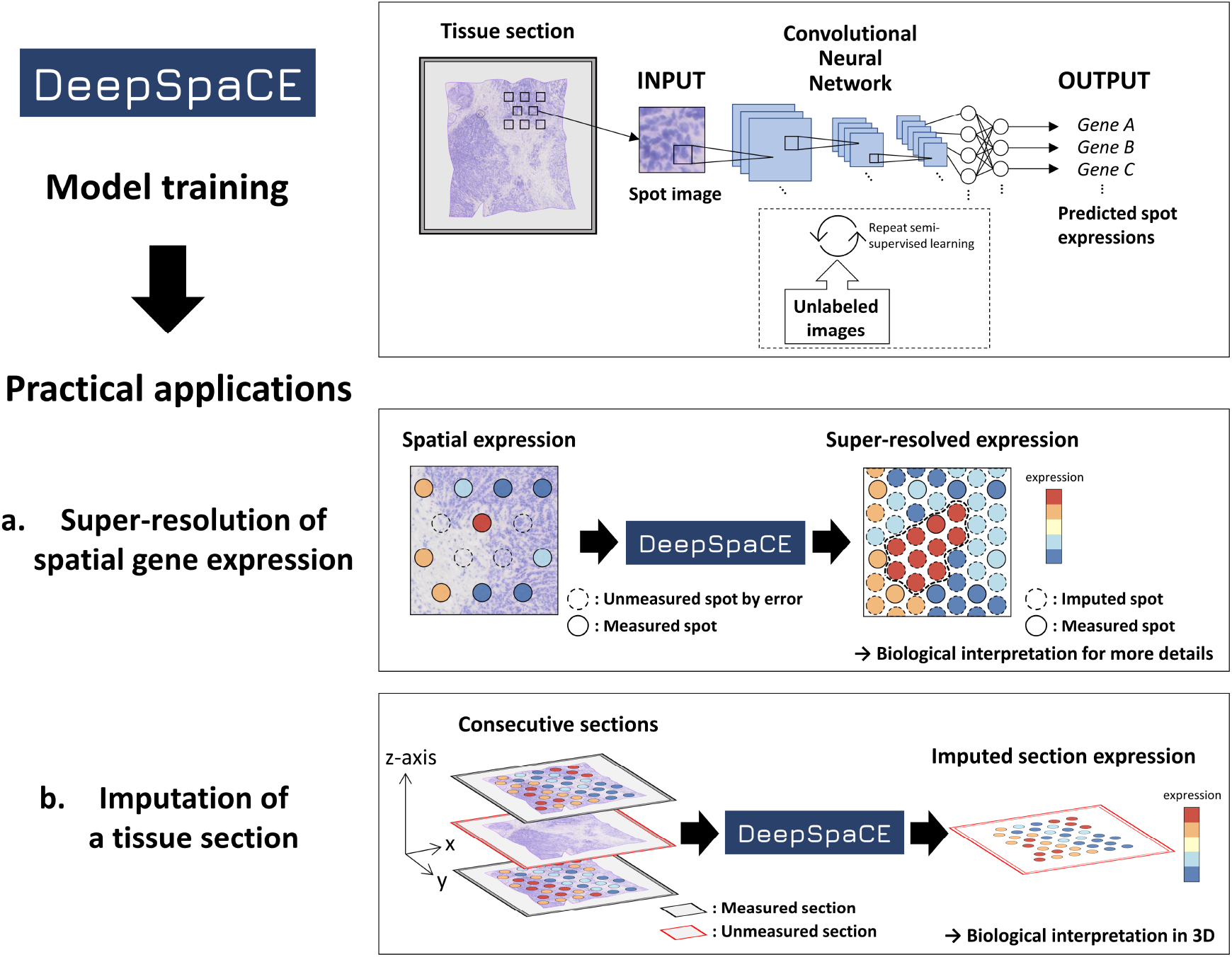
Overview of DeepSpaCE. <u>Deep</u> learning model for <u>Sp</u>atial gene <u>C</u>lusters and <u>E</u>xpression (DeepSpaCE) is a method for predicting gene-expression levels and transcriptomic cluster types from tissue spot images. DeepSpaCE is composed of two parts: the model training part and gene-prediction part. In the case of using semi-supervised learning as an option, unlabeled images are used to improve the prediction accuracy with predicted proxy labels. As practical applications of DeepSpaCE, we conducted super-resolution of spatial gene expression and tissue section imputation. (**a**) Super-resolution was used for predictions with unmeasured spot images (e.g., images among spots whose expression profiles were measured using the *in situ* capturing platform or images on spots with technical errors). Left spatial expression pattern shows that some spots are lacks of expression value because of a technical problem such as permeabilization error (dotted circle). Right image shows an additional spatial expression pattern imputed by DeepSpaCE, and its highly expressed region in the center of the section (dotted line). It is challenging to infer a functional boundary such as cancer infiltration from spatial expression profiles of sparse spots (left). Spatial expression profiles of dense spots imputed by DeepSpaCE and their gene annotations enable to delineate a functional boundary clearly. (**b**) Tissue section imputation was used to predict gene-expression levels in one of tissue section within consecutive sections. By using DeepSpaCE, the unmeasured spatial expression profiles of the slide (red frame) can be imputed by at least one adjacent slide (black frame) whose expression profiles were measured using the *in situ* capturing platform.

Super-resolution is used to predict unmeasured spots in the same image (e.g., images among spots whose expression profiles were measured using the *in situ* capturing platform or images on spots with section-permeabilization errors). Tissue section imputation is performed to predict spatial expression profiles of a section from a series of directly measured consecutive sections. These two applications are helpful for reducing experimental costs and clarifying biological functions at higher resolution and in three dimensions. Because substantially fewer labeled spots are available compared with general deep CNN datasets, we implemented the semi-supervised technique in DeepSpaCE to increase prediction accuracy, as described in detail below. All DeepSpaCE codes for Visium, a standardized, commercially available platform for spatial transcriptome, will be available after acceptance on the GitHub repository (https://github.com/tmonjo/DeepSpaCE).

### Preprocessing of spatial expression data

We preprocessed the spatial expression data from three human breast cancer tissue sections (sections A–C) and their consecutive sections (sections D1–D3) (**Supplementary Fig. S1**). We excluded spots containing few expressed (or measured) genes to filter out spots with potential permeabilization errors, and normalized the spatial expression data to improve the training efficiency by reducing noise. Particularly, the filtering step was critical because spatial expression profiling requires very practiced skills for handling tissue slides and treating reagents homogeneously, and few expressed genes may reveal permeabilization errors in the spots. Indeed, in our spatial transcriptome datasets of human breast cancer tissues, the right bottom regions in sections D1 and D3, as well as the right upper region in section D2, showed undetected unique molecular identifiers (UMIs), indicating that section-permeabilization errors occurred in these regions (**Supplementary Fig. S2a, b**). Similarly, such undetected regions were observed in the 10x Genomics Visium demo data of human heart tissue (**Supplementary Fig. S2c**). We used these undetected spots to evaluate the performance of section imputation by DeepSpaCE, as shown below.

### Prediction and experimental validation of gene-expression profiles and cluster types

We trained the DeepSpaCE models of three breast cancer-marker genes, estrogen receptor 1 (*ESR1*), erb-b2 receptor tyrosine kinase 2 (*ERBB2*), and marker of proliferation Ki-67 (*MKI67*)^13,14^, which were not used during the parameter-optimization procedures (see Methods). We performed the 5-fold cross-validation using section D2 as both a training and a testing set. The Pearson’s correlation coefficients between the measured and predicted values were 0.588 (standard deviation [SD] = 0.025; *ESR1*), 0.424 (SD = 0.050; *ERBB2*), and 0.219 (SD = 0.041; *MKI67*) (**Supplementary Fig. S3**). Notably, comparison of *ESR1* levels in the D2 section by H&E staining highlighted undetected highly expressed spots in the upper right region of section D2, possibly because of a permeabilization error in the Visium experiment (**Fig. 2a**). This was further confirmed by determining the protein-expression pattern observed by immunohistochemical staining of the adjacent section using an *ESR1* antibody (**Fig. 2b**), which was consistent with the predicted expression levels, suggesting the applicability of our DeepSpaCE method for section imputation.

**Figure 2.**
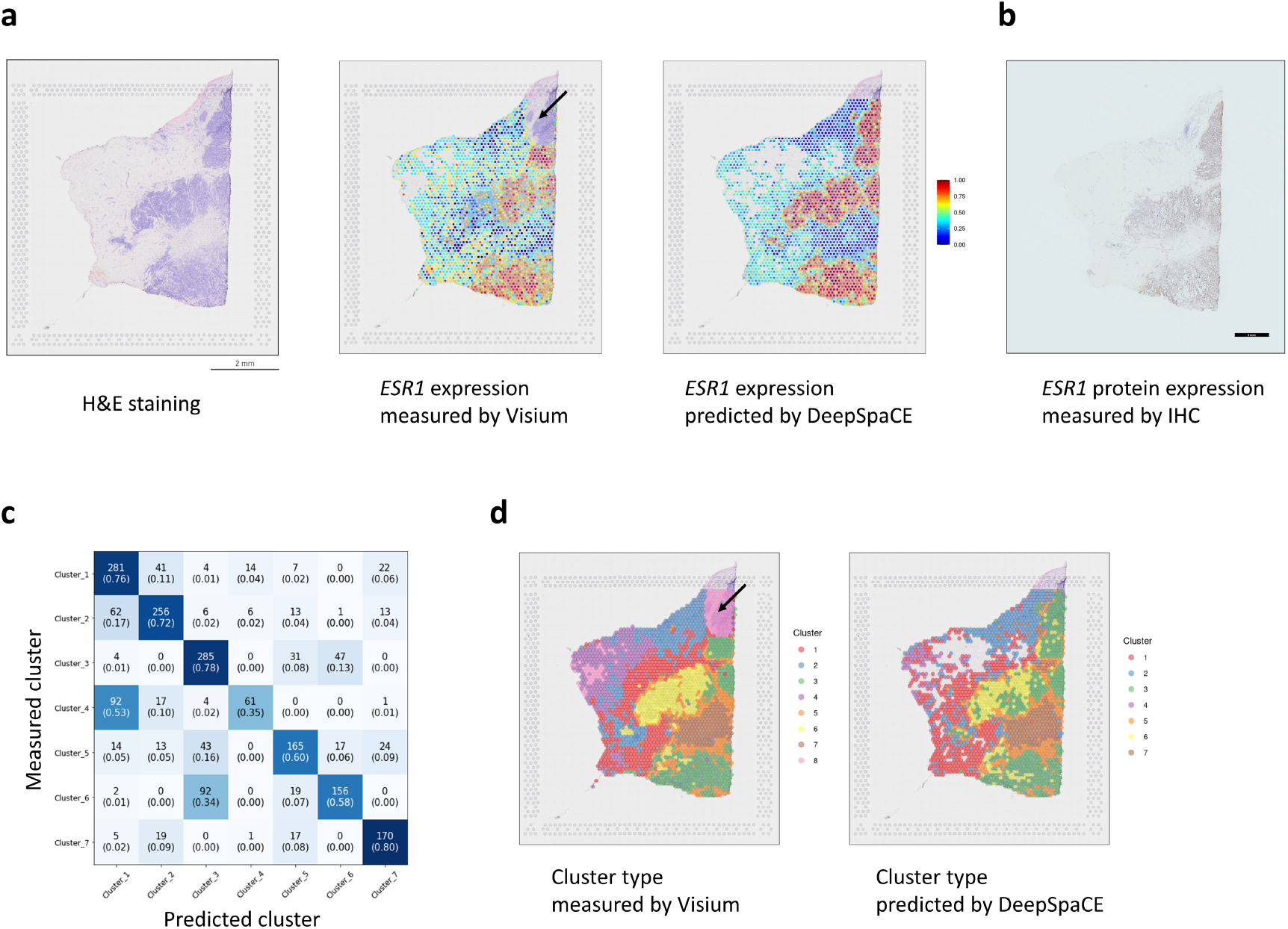
DeepSpaCE predicts spatial gene expression and cluster types. (**a**) Left image shows section D2 after hematoxylin and eosin (H&E) staining. Middle image shows a heatmap of normalized *ESR1* expression in section D2, measured using the 10x Genomics Visium platform. *ESR1* expression in the upper right region (black arrow) of section D2 could not be measured because of permeabilization errors. Right image shows the heatmap of *ESR1* expression in section D2, predicted by DeepSpaCE. The blank areas represent spots that were excluded because of a small amount of information. (**b**) Image showing immunohistochemical staining of ESR1 protein in the adjacent section of section D2. (**c**) Heatmap shows the confusion matrix of the measured and predicted cluster types. The number in each box is the number of labels, and the number inside each pair of parentheses is the recall value. (**d**) Left image shows a heatmap of transcriptomic cluster types in section D2, as measured using the Visium platform. The cluster types in the upper right region (black arrow) of section D2 were not determined because of permeabilization errors. Right image shows the heatmap of cluster types in section D2, predicted by DeepSpaCE. Blank areas represent spots that were excluded because of low information. For training and gene prediction parts, we excluded cluster 8 because most of the region belonging to the cluster showed permeabilization errors.

Next, based on the 5-fold cross-validation in section D2, we assessed the prediction accuracy of transcriptomic cluster type derived from Space Ranger software. By comparing the clusters from Space Ranger with the predicted clusters, we calculated the recall value (see Methods) of the clusters, which ranged from 35% (cluster 4) to 80% (cluster 7) (**Fig. 2c**). Briefly, although the prediction accuracy was low when comparing non-cancerous regions (e.g., clusters 1 and 4), the cluster types between cancer sites and non-cancer sites were clearly distinguishable (e.g., clusters 1 and 3). Similar to the findings described in the previous paragraph, the cluster type was predicted in the unmeasured upper right region of section D2, which showed a permeabilization error (**Fig. 2d**). The predicted types of clusters in this region were plausible based on the spatial transcriptome and DeepSpaCE analysis using the adjacent sections D1 and D3, which could measure the region (**Supplementary Fig. S4**).

### Super-resolution of spatial gene expression

We performed super-resolution for *ESR1* in the images of spots measured in section C (**Supplementary Fig. S5a, b**). Section C was used as both a training and test set to generate a super-resolved image as an example. The super-resolved image of *ESR1* expression in section C was consistent with the results of immunohistochemical staining, supporting that DeepSpaCE enables accurate high-resolution observations of expression profiles (**Supplementary Fig. S5c**). Notably, we found a region with low *ESR1* expressions in the super-resolved image, which was not clearly observed in the original spatial transcriptome datasets (**Supplementary Fig. S5b**), confirming the importance of super-resolution.

We focused on secreted protein acidic and cysteine rich (*SPARC*), a potential cancer-invasion marker, and assessed whether the super-resolution method could facilitate biological interpretations provided by pathologists based on histology patterns in H&E-stained images. After training and validating the DeepSpaCE model (**Supplementary Fig. S6**), we predicted *SPARC*-expression levels among the original spots in section D2 (**Fig. 3a, b**). Although the color tones themselves did not explain the *SPARC* expression patterns in the H&E-stained images, the patterns were successfully predicted by the DeepSpaCE model. Comparison of the super-resolved image of *SPARC* with the H&E-stained image showed that the invasive tumor region overlapped substantially with the distribution of *SPARC* expression (**Fig. 3c**), whereas such patterns were not apparent from the color tones in the original spatial-transcriptome data. *SPARC* is secreted into the extracellular matrix from cancer and stromal cells, and high *SPARC*-mRNA expression is related to metastasis and poor prognosis in several types of cancers^15^. Thus, the super-resolved *SPARC*-expression image highlighted the potential tumor-invasion region and made it easier to identify by a non-pathologist in cases where a given transcript’s function is known.

**Figure 3.**
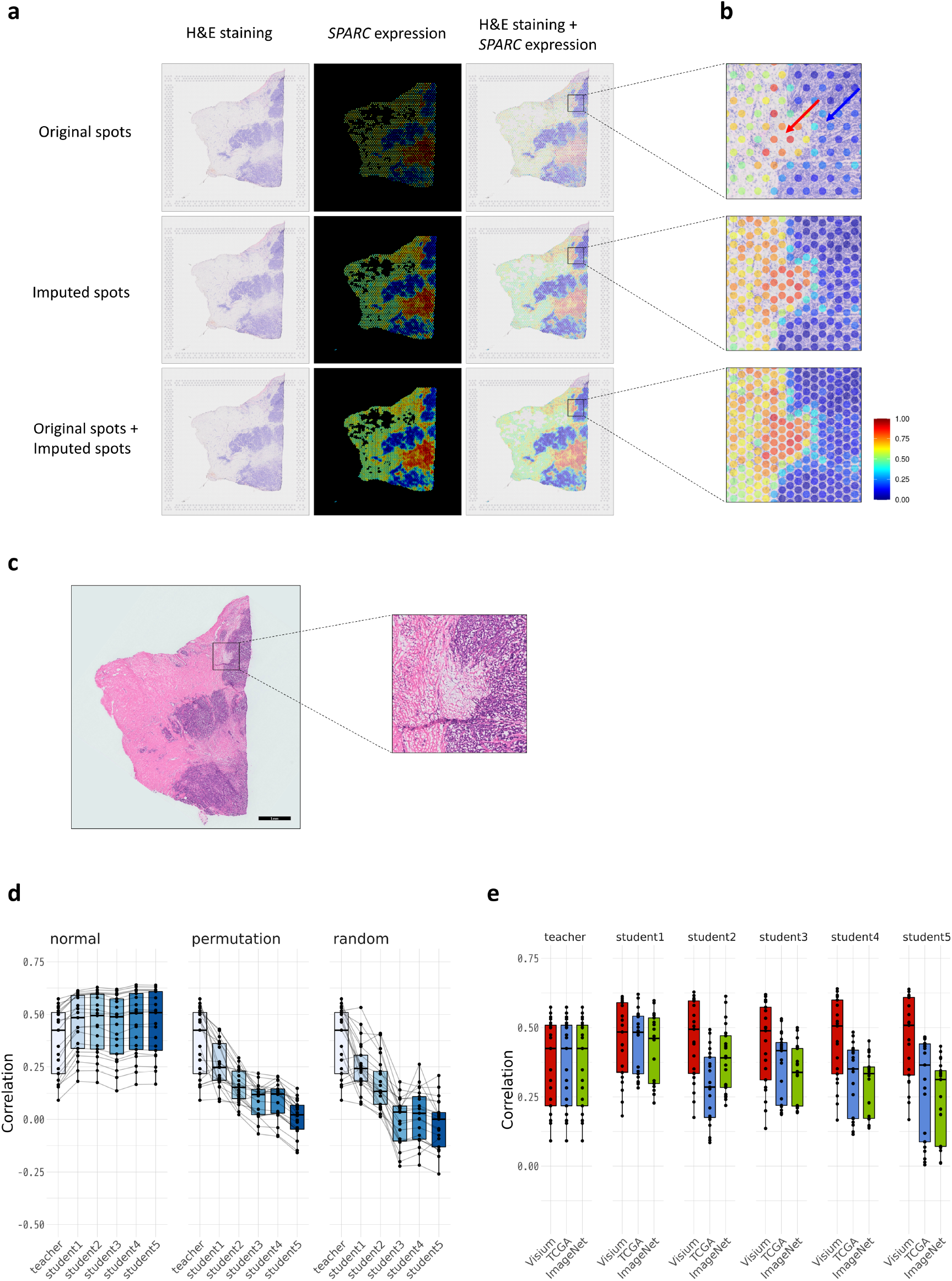
Super-resolution and section imputation as practical applications of DeepSpaCE. Super-resolution of *SPARC* expression using DeepSpaCE highlights tumor invasion more clearly and semi-supervised learning for tissue section imputation using DeepSpaCE improves prediction accuracy. (**a**) Nine images show the super-resolved results for *SPARC* expression. Three images in the left column show section D2 after H&E staining. Three images in the middle column show the heatmaps of predicted *SPARC* expression by DeepSpaCE for the original spots (top), imputed spots (middle), and both original and imputed spots (bottom). Three images in the right column show overlays of predicted *SPARC* expression by DeepSpaCE and H&E staining for section D2. (**b**) Three enlarged images on the right area show tumor cell invasion (blue arrow) and the microenvironment (red arrow). Spot size is adjusted to smaller than the exact spot size of the Visium platform to show the background image. (**c**) Left image shows the H&E-stained section adjacent to section D2. Right enlarged image is the same region as **Fig.3b**. Enlarged image shows the invasion of tumor cells. (**d**) Box plots show Pearson’s correlation coefficients between the measured and predicted gene-expression levels of 21 breast cancer-related microenvironment markers. Left box plot displays the results of semi-supervised learning, which showed increasing Pearson’s correlation coefficients. Middle and right box plots show the semi-supervised learning results with permutated and randomized values. For the box plot, the box indicates the first and third quartiles; horizontal center line marks the medians; upper whisker extends from the hinge to the highest value that is within 1.5 × interquartile range (IQR) of the hinge; lower whisker extends from the hinge to the lowest value within 1.5 × IQR of the hinge; and data were plotted as points. Black lines between boxes connect the same gene. (**e**) Box plots show Pearson’s correlation coefficients between the measured and predicted gene-expression levels of 21 breast cancer-related microenvironment markers. Three types of image sets were compared for semi-supervised learning, namely sections A–C (red); data from The Cancer Genome Atlas (TCGA) (blue); and ImageNet data (green). For the box plot, the box indicates the first and third quartiles; horizontal center line marks the medians; upper whisker extends from the hinge to the highest value that is within 1.5 × IQR of the hinge; lower whisker extends from the hinge to the lowest value within 1.5 × IQR of the hinge; and data were plotted as points.

### Imputation of a tissue section using semi-supervised learning

To assess whether semi-supervised learning can improve the prediction accuracy of DeepSpaCE, we performed tissue section imputation for section D2 using the model trained by sections D1 and D3 (randomly selected 80% spots were used as training data and others were used as validation data). After training the teacher model (first trained model), we developed student models 1–5 by performing five rounds of semi-supervised learning using breast cancer section images (section A– C) as unlabeled images. We selected the most predictive student model in the validation data as the best model. We found an increasing trend in the Pearson’s correlation coefficients between the measured and predicted expression levels, as expected (**Supplementary Table S1**). In the teacher model, the mean Pearson’s correlation coefficient for 21 genes was 0.369. The mean Pearson’s correlation coefficients progressively increased for student model 1 (0.414), student model 2 (0.455), student model 3 (0.438), student model 4 (0.457), and student model 5 (0.458) (**Fig. 3d**). For *SPARC*, Pearson’s correlation coefficient increased from 0.509 (SD = 0.069; teacher model) to 0.616 (SD = 0.067; student model 4). For *MXRA5*, the Pearson’s correlation coefficient was not increased after analysis using student model 1, although it was increased from 0.501 (SD = 0.083; teacher model) to 0.527 (SD = 0.093; student model 1). Thus, the semi-supervised learning method may increase the accuracy of DeepSpaCE through additional computational costs.

To verify whether their related unlabeled images could improve the accuracy of the DeepSpaCE model, we performed semi-supervised learning with permutated gene-expression levels or randomized the values as negative controls. The Pearson’s correlation coefficients did not increase, but rather decreased, when permutated or randomized values were used. Moreover, Pearson’s correlation coefficients did not increase when irrelevant images of dogs or cats (obtained from ImageNet) were used for semi-supervised learning. Furthermore, Pearson’s correlation coefficients did not increase when breast cancer section images obtained from The Cancer Genome Atlas (TCGA) were used for semi-supervised learning (**Fig. 3e**).

## Discussion

In this study, we proposed performing super-resolution and section imputation with DeepSpaCE and validated the accuracy from cross-validation and immunohistostaining. These approaches made it possible to derive more knowledge from existing spatial transcriptome datasets. As a compelling example, the relationship between *SPARC* expression and cancer invasion was highlighted via super-resolution, whereas detecting the invasive region using original spatial transcriptome data was difficult because the measured spots were not dense (**Fig. 3a, b**). The SPARC glycoprotein has a high affinity for albumin, and macrophage-derived SPARC contributes to metastasis by acting at the step of integrin-mediated migration of invasive cells^15^. Previously, *SPARC* mRNA expression was reported as a predictor of a pathological complete response after neoadjuvant nab-paclitaxel therapy^16^. Our study underscored the relationship between *SPARC* expression and invasive regions, which may be clinically important for treating breast cancer. This interpretation does not require expertise in histology or pathology but requires gene annotations, with should be familiar to researchers of spatial transcriptome. In addition, super-resolution in section C identified the region with low expressed *ESR1*, the amplification of which is frequently observed in proliferative breast cancers^13^. Although it is unclear whether the region indicates the heterogeneity of breast cancer tissues or existence of normal tissues, this region was unclear in the original spatial transcriptome data and expression in adjacent sections was experimentally validated.

For super-resolution and section imputation, we developed DeepSpaCE to predict expression levels from spot images from Visium. This spot-level analysis should reveal more detailed patterns than those obtained by CNN using pairs of images of the bulk transcriptome^17^ by resolving spatial expression patterns. DeepSpaCE requires a minimum of a single experiment to analyze the spatial transcriptome; nevertheless, the predictions were well-validated by cross-validation and experimental analysis. DeepSpaCE as well as super-resolution and section imputation methods aim to maximize the value of existing datasets and provide foundations for subsequent experiments from at least a single dataset without additional experimental costs. This is an important difference from the recently proposed STNet study in which trained spatial transcriptome data (not from the Visium platform) was obtained from as many as 23 individuals^18^.

The number of training datasets used for single spatial transcriptome analysis (maximum 4,992 spots/slide with the Visium platform) was not sufficient for training the CNN in general, as a previous study used ∼557,000 images from 830 slides to predict lung cancer subtypes and ∼212,000 images from ∼320 slides to predict lung cancer gene mutations^11^. To increase the ability to apply DeepSpaCE to many datasets for which it is challenging to train the connections between H&E-stained images and expression levels, we implemented a semi-supervised learning method^19^ in DeepSpaCE. The DeepSpaCE model with semi-supervised learning using sections A–C as unlabeled images showed better performance than a simple prediction model using only experimentally obtained spatial transcriptome data. Although we increased the predictive accuracy of tissue section imputation in this case, the Pearson’s correlation coefficients were not improved when using breast cancer H&E-stained images obtained from TCGA as unlabeled images. This may be because the DeepSpaCE model is sensitive to the protocol for obtaining the H&E images (i.e., batch effects disturb the training steps). Therefore, the model that gives the performs the best prediction accuracy when using semi-supervised learning as an option should be determined. In conclusion, DeepSpaCE is an all-in-one package that augments spatial transcriptome data obtained from the *in situ* capturing platform; its applications can improve the understanding of histological expression profiles.

## Methods

### Ethical approval

Breast tissue samples and relevant clinical data were obtained from patients undergoing surgery at St. Marianna University School of Medicine Hospital after obtaining approval from the Clinical Ethics Committee of St. Marianna University (approval number: 2297-i103). The approval allowed the retrieval of surgical pathology tissues that were obtained with informed consented or that were approved for use with a waiver of consent.

### Spatial-transcriptomics datasets

We used six human breast cancer tissue sections, including sections A–C and consecutive sections D1–D3, which were derived from one patient. The spatial transcriptomics experiments were conducted with the same protocol reported in Nagasawa et al.^8^. Briefly, the tissue sections were stained with H&E, and TIFF images were obtained using a microscope at 10× magnification. Spatial-transcriptome profiling was performed using the Visium platform with the standard protocol provided by 10x Genomics (Pleasanton, CA, USA). UMI counts were calculated using 10x Genomics Space Ranger software (version 1.0.0). Visium demo data (version 1.0.0) for the human heart tissue was obtained from the 10x Genomics website (https://www.10xgenomics.com/resources/datasets/).

### Preprocessing of spatial gene-expression data

Regarding the spatial-transcriptome profiles obtained from the Space Ranger pipeline (10x Genomics), we removed spots with low total UMI counts (<1,000) or a low number of measured genes (<1,000). The SCTransform function of Seurat package (version 3.1.4)^20^ was applied to normalize the UMI counts, based on regularized negative binomial regression^21^. Min-max scaling was performed to adjust the expression values between zero and one. We trained 24 genes including three breast cancer-marker genes (*MKI67, ESR1, ERBB2*) and 21 breast cancer-related microenvironment marker genes (*SPARC, IFI27, COL10A1, COL1A2, COL3A1, COL5A2, FN1, POSTN, CTHRC1, COL1A1, THBS2, PDGFRL, COL8A1, SULF1, MMP14, ISG15, IL32, MXRA5, LUM, DPYSL3*, and *CTSK*). These 21 genes were manually selected from the cluster of genes overexpressed in the breast cancer-related microenvironment region. These two gene sets of three genes and 21 genes were respectively used in the training part of DeepSpaCE. The graph-based clustering algorithm^22^ implemented in Space Ranger was used for transcriptomic cluster type prediction.

### Preprocessing of tissue section images

Each spot image was cropped from a tissue slide image, based on the position table in the Space Ranger outputs (**Supplementary Table S2**). We filtered out whitish images in which more than half of the pixels were the >80% percentiles of mean RGB values, as shown below. For image augmentation, we randomly applied image-transform functions of flipping (RandomRotate90, Flip, and Transpose), cropping (RandomResizedCrop), noise (IAAAdditiveGaussianNoise and GaussNoise), blurring (MotionBlur, MedianBlur, and Blur), distortion(OpticalDistortion, GridDistortion, IAAPiecewiseAffine, and ShiftScaleRotate), contrast (RandomContrast, RandomGamma, and RandomBrightness), and color-shifting (HueSaturationValue, ChannelShuffle, and RGBShift) in Albumentations library (version 0.4.5)^23^.

### Preprocessing of images obtained from TCGA and ImageNet

We obtained 1,978 images of H&E-stained TCGA breast cancer sections from the GDC Data Portal (https://portal.gdc.cancer.gov) on August 05, 2020. As negative controls, we obtained 14,500 irrelevant images such as dogs and cats (n02106662, n02110341, n02116738, n02123045, n02123159, n02123394, n02123597, n02124075, n02497673, and n03218198) from ImageNet (http://www.image-net.org) on October 09, 2020. All images obtained from TCGA and ImageNet were cropped to 224 × 224 pixels (**Supplementary Fig. S7**). Four thousand cropped images were randomly selected as unlabeled images for each semi-supervised learning model.

### Training and prediction of gene-expression profiles and transcriptomic cluster types

All deep-learning models were implemented using deep-learning framework PyTorch (version 1.5.1)^24^. We adapted the VGG16 architecture for deep CNN model that has 16 weight layers^25^. We modified the number of output features in VGG16 from 1,000 to the number of genes or cluster types. We simultaneously trained multi genes such as three genes of breast cancer markers or 21 breast cancer-related microenvironment markers. For transcriptomic cluster type predictions, the loss value of the training DeepSpaCE dataset was calculated using the CrossEntropyLoss function. For gene-expression predictions, the loss value was determined as the sum of loss calculated with the SmoothL1Loss function for each gene. As an optimizer, we used Adam^26^ with the hyperparameters of learning rate: 1e-4 and weight decay: 1e-4. Each training was repeated for 50 epochs to stabilize the loss curves (**Supplementary Fig. S8**). Early stopping was applied if the loss value for the validation data did not decrease over five continuous epochs. To evaluate the accuracy of cluster type prediction, we used the recall value which reflects the proportion of positives identified correctly among the actual number of positives (recall = true positive / (true positive + False negative)). For section D2, cluster 8 was excluded from the training set because it consists of a region of permeabilization errors.

### Parameter optimization of DeepSpaCE

We optimized the parameters of DeepSpaCE, such as the image size, image-filtering threshold, and image-augmentation methods. We performed 5-fold cross-validation using six sections (A–C, and D1–D3) to evaluate the prediction accuracy. We developed prediction models for the expression levels of the 21 breast cancer-related microenvironment marker genes (described above) because these genes are representative markers of heterogeneous ductal carcinoma tissues. First, we assessed the impact of the size of the input images (0%, 50%, 100%, 150%, and 200%; relative to the original spot image size) on the prediction accuracies; the results showed that an image size of 150% gave better outcomes than the original and smaller image sizes (**Supplementary Fig. S9a**).

This result is biologically plausible because the surrounding cells can communicate with cells in the spot and affect their gene-expression levels. Second, we assessed the different image-filtering thresholds to exclude uninformative images (i.e., excluding almost white images). We calculated whiteness for each spot by calculating the mean RGB values and obtained the percentiles (50%, 60%, 70%, 80%, 90%, and 100%) over spots in a slide. We filtered out images in which more than half of the pixels were the >80% percentiles of mean RGB values as judged from the histogram (**Supplementary Fig. S9b**). This strategy maximized the prediction accuracy (**Supplementary Fig. S9c**). Third, to further improve accuracy, we augmented images with various image transformations such as flipping, cropping, blurring, distortion, noise, contrast, and color-shifting (**Supplementary Fig. S10**). All image augmentation (except for color-shifting) improved the Pearson’s correlation coefficients compared with using non-augmented images (**Supplementary Fig. S9d**); however, we also used the color-shifting method because H&E-staining on different slides may change the color tones.

### Super-resolution of spatial gene expression

To impute the expression levels among spots on a slide image, new spot image files were created by cropping around three adjacent spots (**Supplementary Fig. S11**). We used sections C and D2 as both the training and test sets (randomly selected 80% spots were used as training data and others were used as test data). By performing super-resolution, the numbers of spots increased from 2,238 to 6,733 and from 2,168 to 6,623 in sections C and D2, respectively. We trained the both of three breast cancer-marker genes and 21 breast cancer-related microenvironment marker genes, respectively. Semi-supervised learning was not used for super-resolution.

### Imputation of a tissue section using semi-supervised learning

Sections D1–D3 were obtained as consecutive sections. Thus, sections D1 and D3 were used as the training set. Section D2 was used as the test set to impute gene-expression levels because it was located between sections D1 and D3. Sections A–C were used for semi-supervised training as unlabeled images. In the noisy student model^27^, gene-expression levels in unlabeled images were predicted using the first trained model (teacher model). Four thousand predicted proxy labels and the associated images were added to the original dataset and used to train the next model, which was designated as a student model. The training student models were run five times (**Supplementary Fig. S12a**). In addition to the spot images of section A–C, we used images of breast cancer sections obtained from TCGA as unlabeled images. Irrelevant images obtained from ImageNet were used as negative controls during semi-supervised learning. In addition, we also performed semi-supervised learning with permuted gene expression and random values as negative controls (**Supplementary Fig. S12b**).

### Immunohistochemistry and measurement of protein expression

Breast cancer tissues were frozen and embedded in optimal cutting temperature compound (Sakura Finetek, Tokyo, Japan). Ten-micrometer-thick sections were cut onto slides using a Leica CM3050 S cryostat (Wetzlar, Germany), fixed in methanol at -20°C for 20 min, and air-dried for 60 min. Endogenous peroxidase activity was blocked in phosphate-buffered saline containing 3% H_2_O_2_ for 5 min. For *ESR1* staining, the sections were incubated with an anti-*ESR1* antibody (FLEX Monoclonal Rabbit Anti-Human Estrogen Receptor α, Agilent technologies, Dako, Glostrup, Denmark) at a 1:2 dilution for 60 min at room temperature. Antibody labeling was detected with the Histofine Simple Stain, MULTI (Nichirei Bioscience, Tokyo, Japan) following the manufacturer’s protocol, and all sections were counterstained with H&E.

### Data processing and analysis

Python (version 3.6.5) was used for preprocessing and implementation of DeepSpaCE with the libraries, torch (version 1.5.1), torchvision (version 0.6.1), numpy (version 1.19.0), pandas (version 1.0.5), scikit-learn (version 0.23.1), mlxtend (version 0.17.2), albumentations (version 0.4.5), opencv-python (version 4.2.0.34), and matplotlib (version 3.2.2). R (version 3.6.0) was used for statistical analysis and visualization with the packages, dplyr (version 1.0.2), data.table (version 1.12.8), Matrix (version 1.2.17), grid (version 3.6.0), rjson (version 0.2.20), hdf5r (version 0.9.7), readbitmap (version 0.1.5), ggplot2 (version 3.3.0), hrbrthemes (version 0.8.0), ggsci (version 2.9), ggpubr (version 0.4.0), cowplot (version 1.0.0), and Seurat (version 3.1.4.9904).

## Code availability

All codes for DeepSpaCE are available on GitHub (https://github.com/tmonjo/DeepSpaCE) (*will be available after acceptance*). These codes include image-preprocessing procedures and expression data produced from Space Ranger.

## Data availability

All sequencing data and pathological images for Visium have been deposited in the DNA Data Bank of Japan under the accession number xxx (*under registration*).

## Acknowledgments

The super-computing resource provided by the Human Genome Center (The University of Tokyo) and AI Bridging Cloud Infrastructure (ABCI), provided by the National Institute of Advanced Industrial Science and Technology (AIST), were used to develop and validate DeepSpaCE.

## Author Contributions

TM, MK, and YK conceived the study and analyzed the data. TM developed and implemented the DeepSpaCE algorithm. SN and YS contributed to the Visium data acquisition. SN contributed to the immunohistochemistry experiments. TM and MK wrote the manuscript with critical input from SN, YS, and YK.

