## Supplementary Information for "Efficient prediction of a spatial transcriptomics profile better characterizes breast cancer tissue sections without costly experimentation"

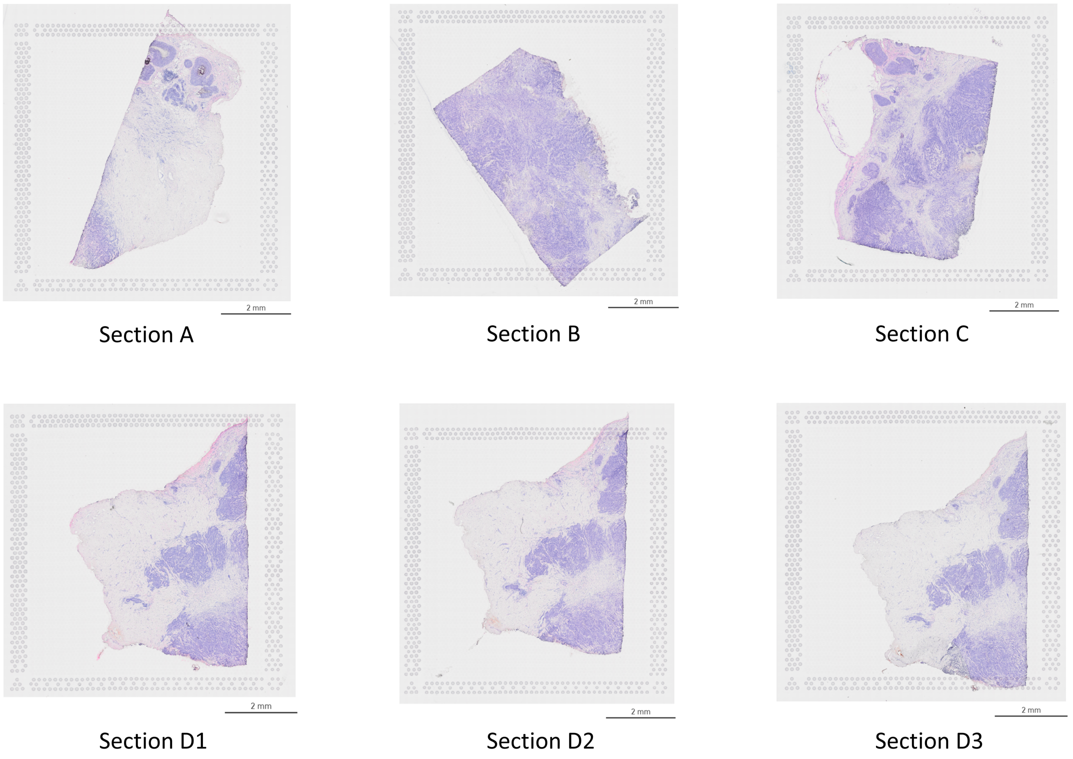


**Supplementary Figure S1** Human breast cancer tissue sections. Six slide images show H&E-stained human breast cancer tissues from sections A–C, and consecutive sections D1–D3. All sections were derived from one patient.

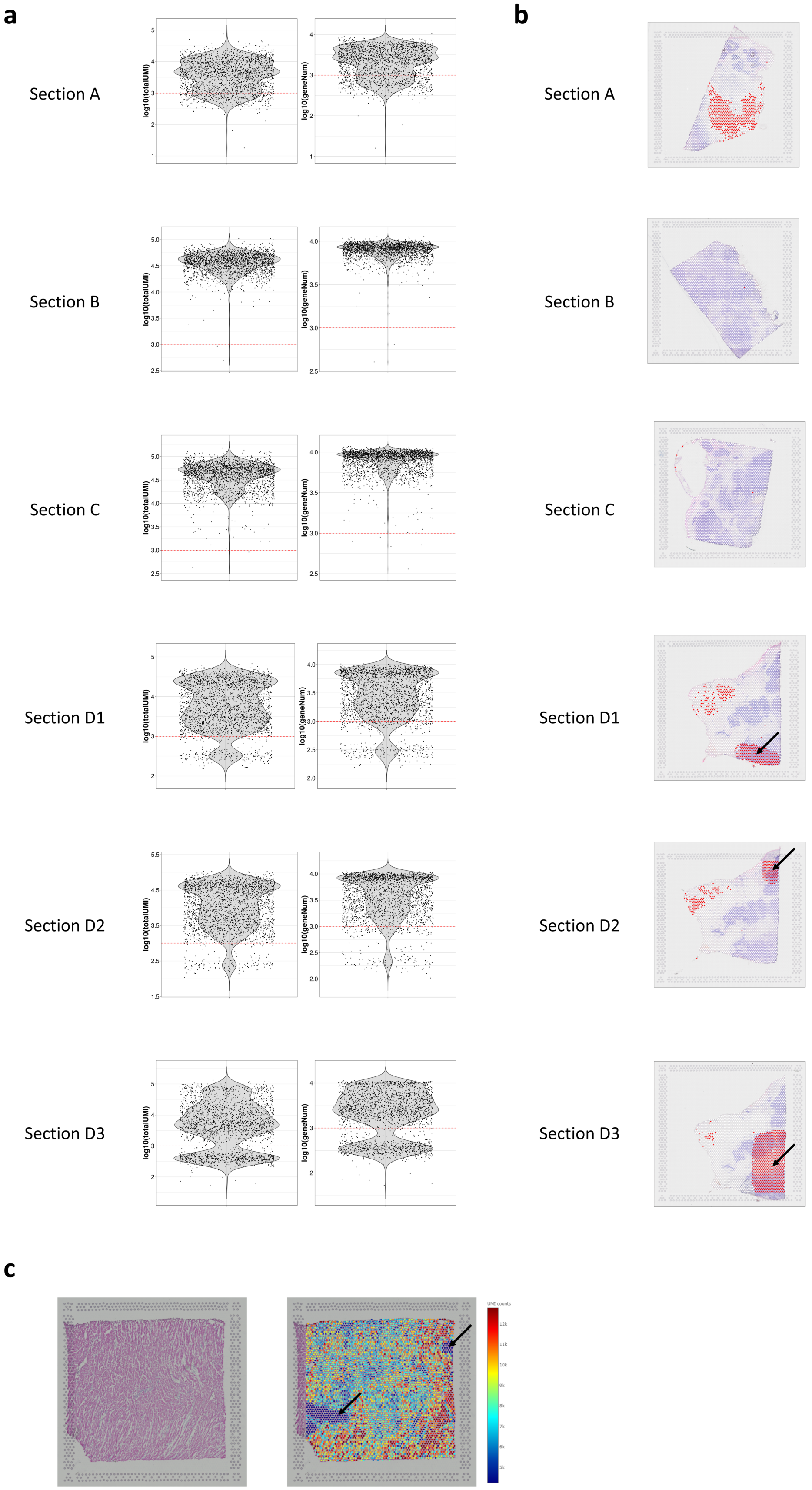
a
b
Section A
Section B
Section C
Section D1
Section D2
Section D3
log10(totalUMI)
log10(geneNum)
c
UMI counts

9

**Supplementary Figure S2** Gene-expression normalization and filtering. (**a**) Violin plot shows the log10 transformed total unique molecular identifier (UMI) counts (left) and number of measured genes (right) in sections A–C and D1–D3. The spots under the red line were filtered out. For the violin plot, the red horizontal line is the threshold of spot filtering, and the data were plotted as points. (**b**) Heatmaps overlaid on H&E-stained sections showed that the red spots were removed because of low UMI counts (or the number of measured genes) in sections A–C and D1–D3. The right bottom regions in sections D1 and D3, as well as the right upper region in section D2, showed undetected regions because of permeabilization error (black arrows) (**c**) Left image shows the H&E-stained section of human heart tissue obtained from 10x Genomics website. The right image shows the heatmap of total UMI counts measured by Visium in the human heart tissue section. The heatmap image was obtained from Space Ranger. Total UMI counts were extremely low in the upper right and the lower left region of the section (black arrows), likely because of a technical problem such as permeabilization error.

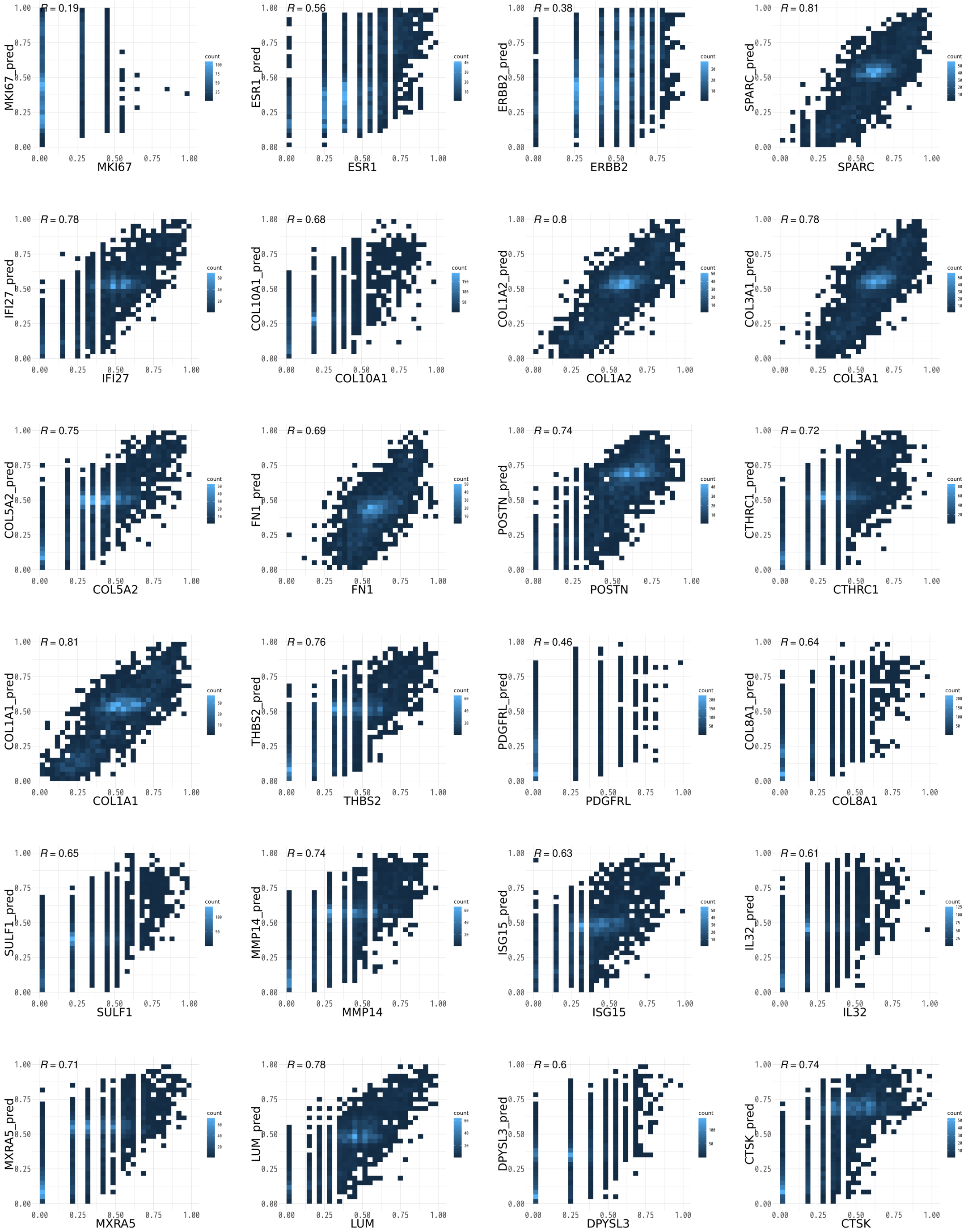


**Supplementary Figure S3** Prediction of gene expression. The plots show the expression values obtained from the 5-fold cross-validation in section D2. 24 plots are including three breast cancer-marker genes (MKI67, ESR1, ERBB2) and 21 breast cancer-related microenvironment marker genes. The x-axis is the measured expression value, and the y-axis is the predicted value. The intensity of the color indicates the number of dots. The upper left value is the Pearson's correlation coefficient between measured and predicted values.

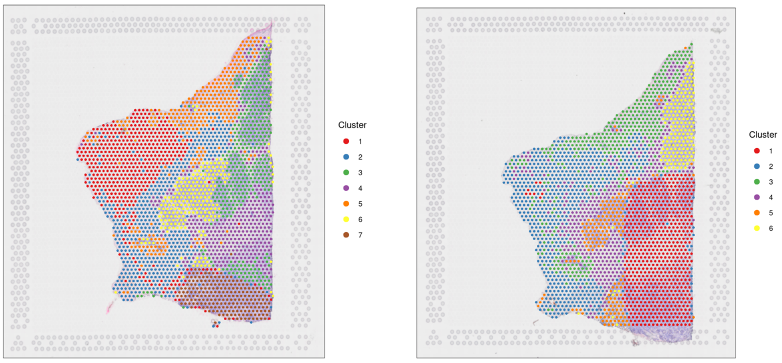


Section D1 Section D3

**Supplementary Figure S4** Transcriptomic cluster types. Image on the left shows the cluster types obtained from Space Ranger in section D1. Bottom right of section D1 was classified as cluster 7 because of permeabilization errors. Right image shows the cluster types obtained from Space Ranger in section D3. Bottom right of section D3 was classified as cluster 1 because of permeabilization errors.

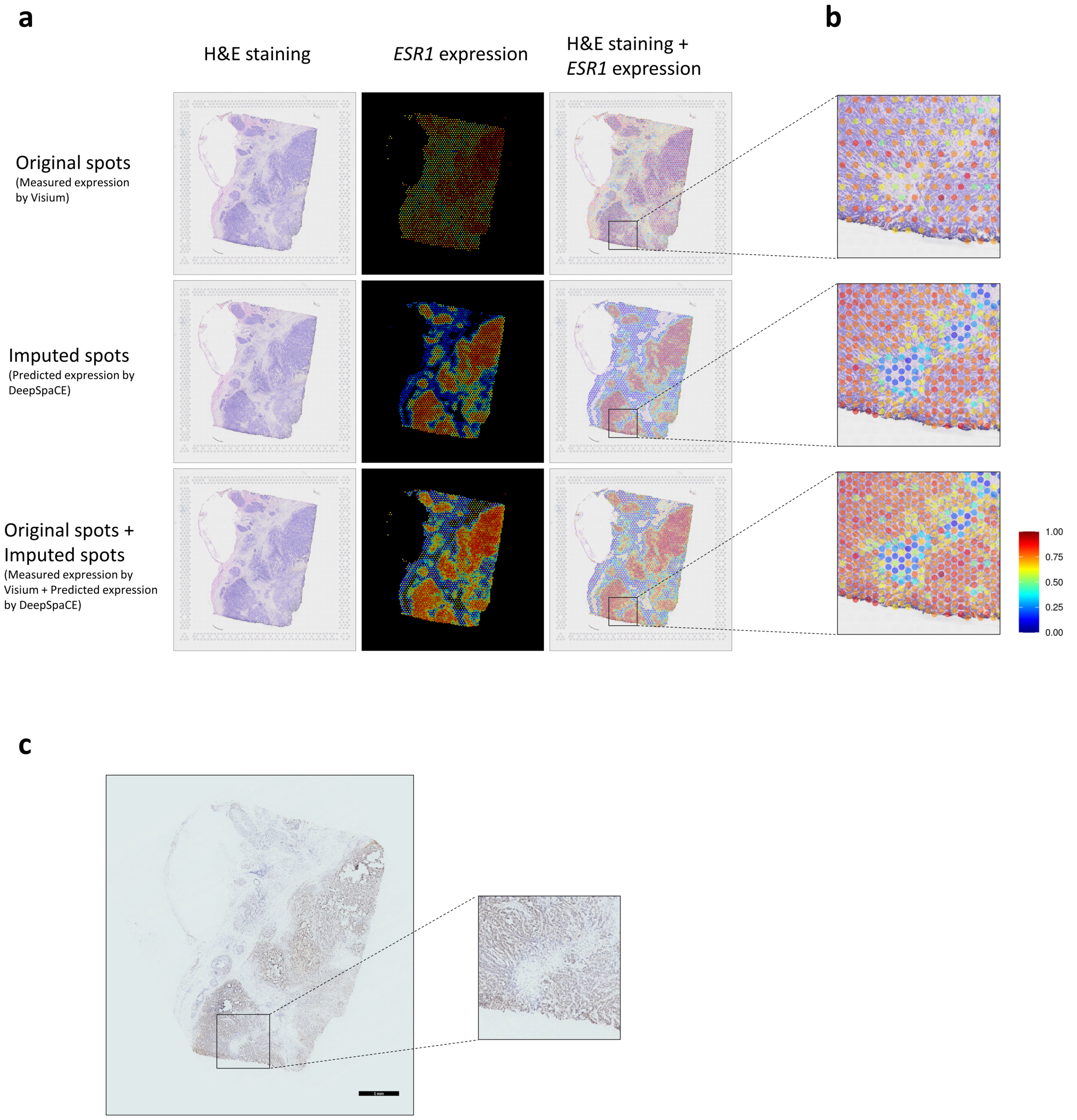


**Supplementary Figure S5** Super-resolution of *ESR1*. (**a**) Nine images show the super-resolved results of *ESR1* expression. The three images in the left column show section C after H&E staining. The three images on the middle column represent heatmaps of *ESR1* expression in the original spots measured by Visium (top), imputed spots predicted by DeepSpaCE (middle), and both original spots measured by Visium and imputed spots predicted by DeepSpaCE (bottom). The three images on the right columns show *ESR1* expression overlaid on section C after H&E staining. (**b**) Three enlarged images show the region of low *ESR1* expressions. Spot size is adjusted to smaller than the exact spot size of the Visium platform to show the background image. (**c**) Image of section C after immunohistochemical staining of the *ESR1* protein. Right enlarged image is the same region as **Supplementary Fig. S5b**.

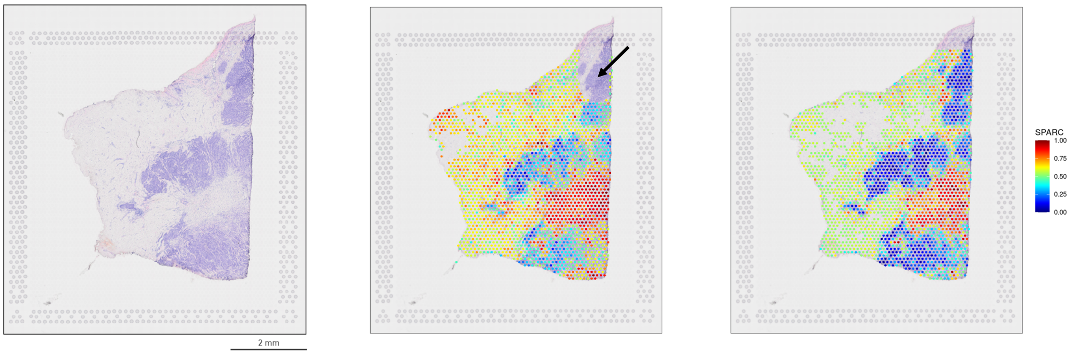


H&E staining

*SPARC* expression measured by Visium

*SPARC* expression predicted by DeepSpaCE

**Supplementary Figure S6** Measured and predicted expression of *SPARC*. Left image shows section D2 after hematoxylin and eosin (H&E) staining. Middle image shows a heatmap of normalized *SPARC* expression in section D2, measured using Visium. *SPARC* expression in the upper right region (black arrow) of section D2 could not be measured, because of permeabilization errors. Right image shows the heatmap of *SPARC* expression in section D2, predicted by DeepSpaCE. Blank areas represent spots that were excluded because of a low amount of information.

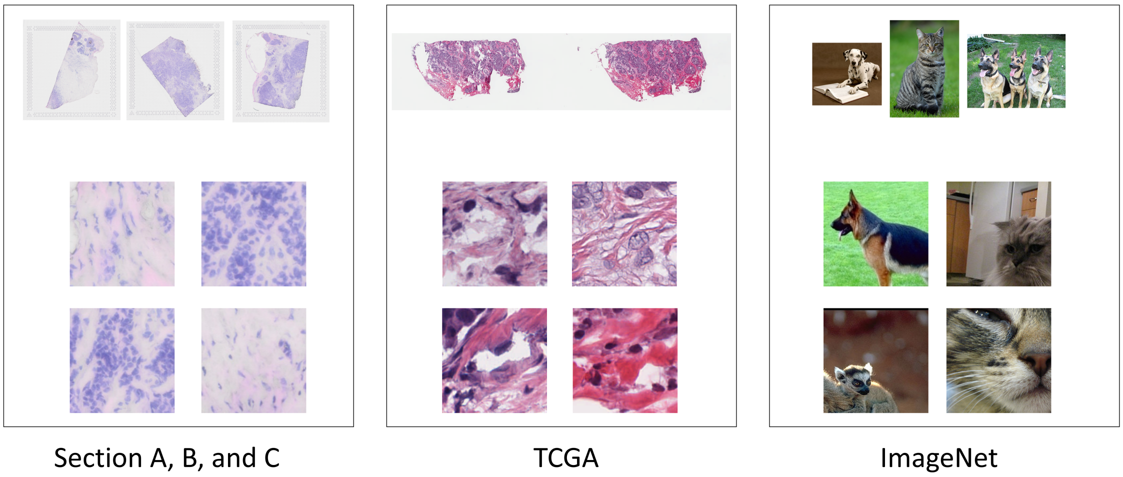


**Supplementary Figure S7** Examples of unlabeled images used for semi-supervised learning. Left images are derived from sections A–C, center images are obtained from TCGA, and right images are obtained from ImageNet. Original images obtained from TCGA and ImageNet were cropped to 224 × 224 pixels.

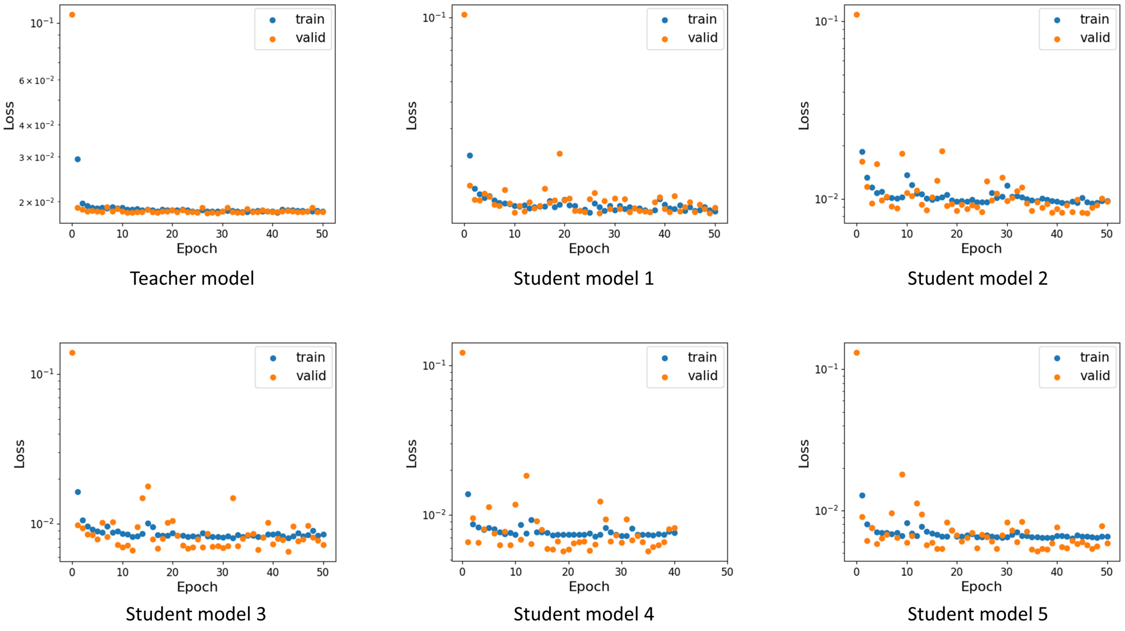


**Supplementary Figure S8** Examples of a learning curve. Scatter plots show the loss values for the training (blue) and validation (orange) datasets. The learning curves were obtained in the semi-supervised learning of 21 breast cancer-related microenvironment marker genes. The training cycle was repeated until 50 epochs. In the case of student model 4, the learning was terminated by early stopping.

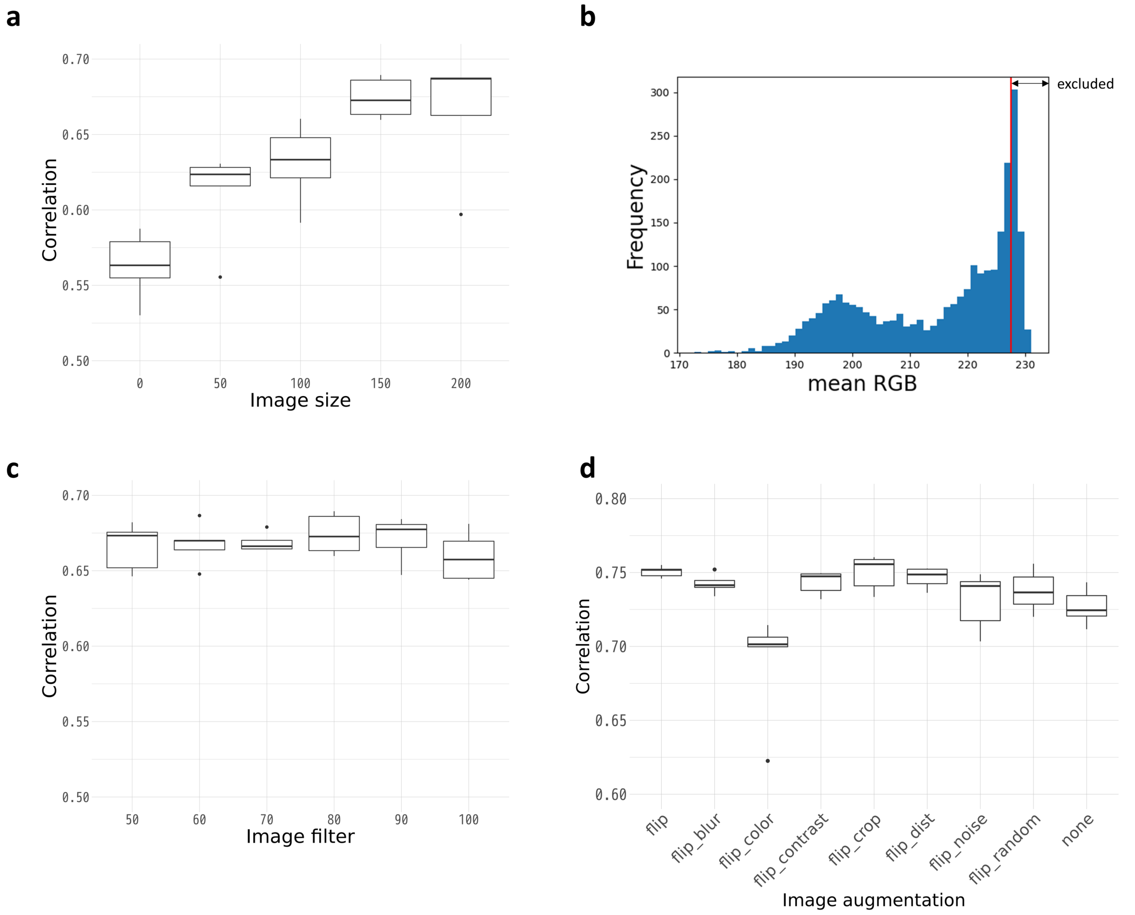


**Supplementary Figure S9** Parameter optimization of DeepSpaCE. (**a**) Box plot shows the Pearson's correlation coefficients for each image size (0%, 50%, 100%, 150%, and 200%; relative to the original Visium spot size). (**b**) Histogram shows the mean RGB values in section D2. Spots on the right of the red line were filtered out. (**c**) Box plot shows Pearson's correlation coefficients for each image-filtering threshold (50%, 60%, 70%, 80%, 90%, and 100%; percentiles of mean RGB values). (**d**) Box plot shows the Pearson's correlation coefficients for each image-augmentation method (flipping, flipping + blurring, flipping + color, flipping + contrast, flipping + cropping, flipping + distortion, flipping + noise, flipping + random (blurring, distortion, or noise), and none). For the box plot, the box indicates the first and third quartiles; the horizontal center line marks the medians; the upper whisker extends from the hinge to the highest value that is within 1.5 × interquartile range (IQR) of the hinge; the lower whisker extends from the hinge to the lowest value within 1.5 × IQR of the hinge.

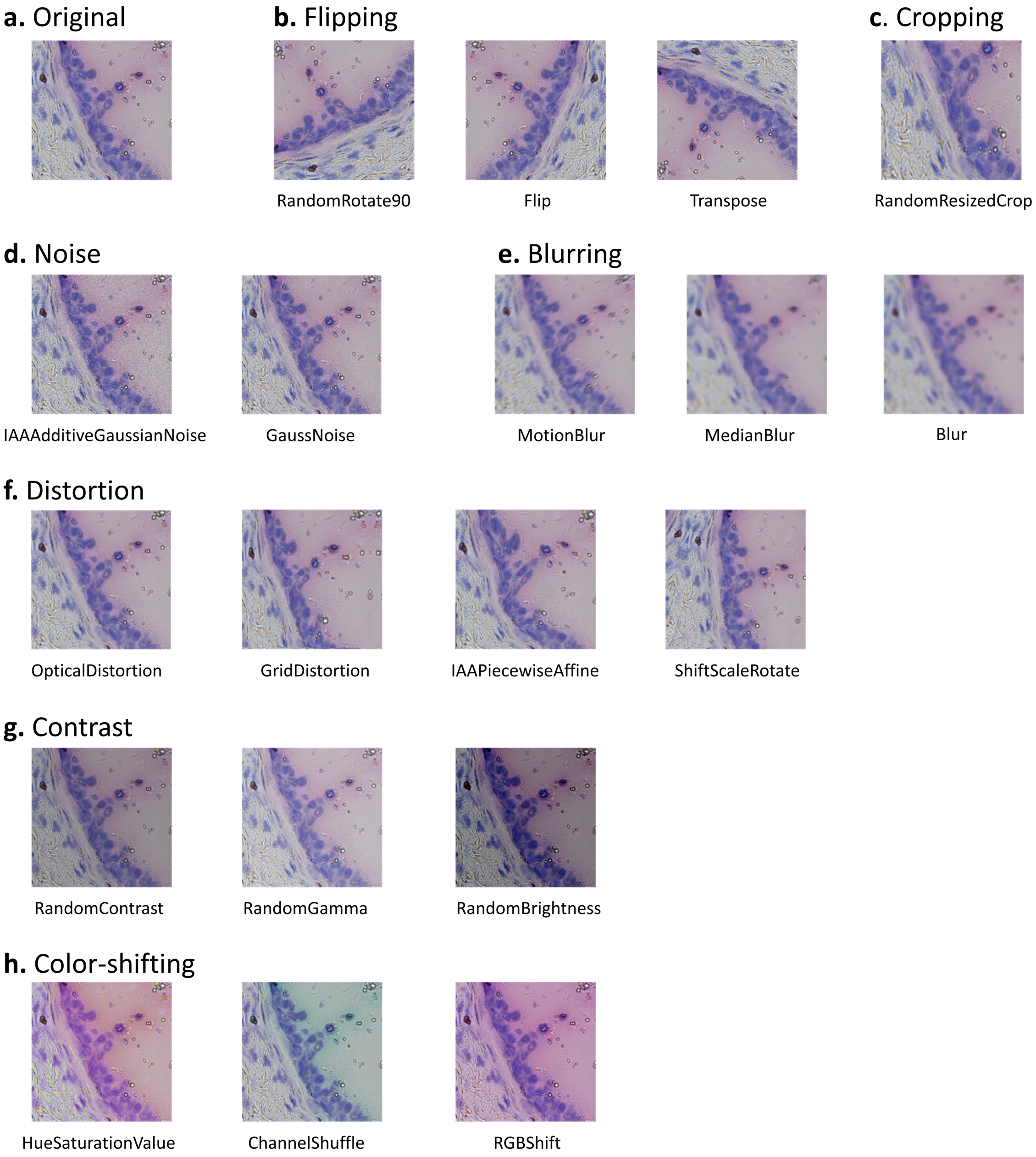


**Supplementary Figure S10** Examples of image augmentation. Examples of each image augmentation (generated using the Albumentations library) are shown. (**a**) Original image. (**b**) Flipping (RandomRotate90, Flip, and Transpose). (**c**) Cropping (RandomResizedCrop). (**d**) Noise (IAAAdditiveGaussianNoise and GaussNoise). (**e**) Blurring (MotionBlur, MedianBlur, and Blur). (**f**) Distortion (OpticalDistortion, GridDistortion, IAAPiecewiseAffine, and ShiftScaleRotate). (**g**) Contrast (RandomContrast, RandomGamma, and RandomBrightness). (**h**) Color-shifting (HueSaturationValue, ChannelShuffle, and RGBShift).

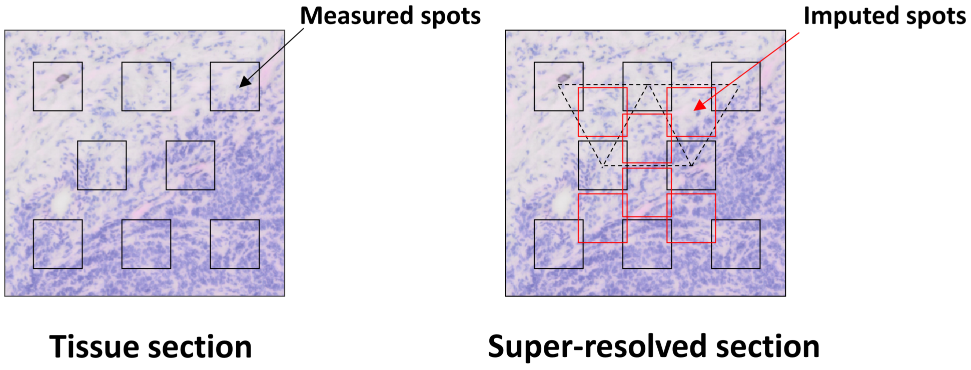


**Supplementary Figure S11** Cropping for super-resolution of section images. Left section image shows the original spots (black) measured by the *in situ* capturing platform. Right section image shows the original spots (black) and imputed spots (red) which is located on the center around three original spots.

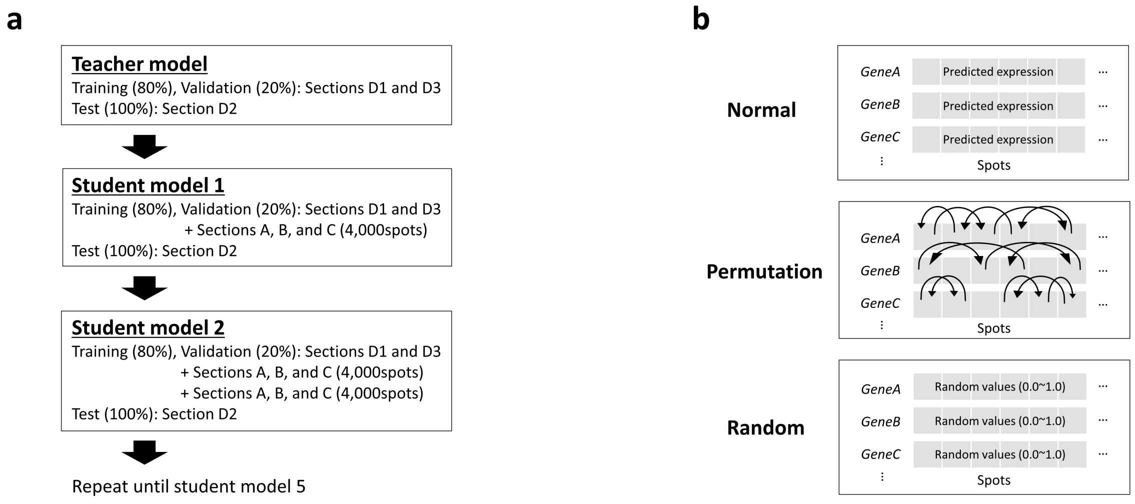


**Supplementary Figure S12** Schematic of semi-supervised learning. (**a**) Schematic of semi-supervised learning showing the dataset for each model in the section imputation. (**b**) We compared three types of gene-expression profiles using semi-supervised methods. In the case of usual values, the predicted expression levels were directly used for semi-supervised learning. In case of permutated values, the predicted expression levels were randomly permutated and used for semi-supervised learning. Random values of zero to one were used for semi-supervised learning.

| Gene symbol | Teacher model | Student model 1 | Student model 2 | Student model 3 | Student model 4 | Student model 5 |
|---|---|---|---|---|---|---|
| **IFI27** | 0.574 (0.030) | 0.599 (0.060) | 0.605 (0.081) | 0.600 (0.067) | 0.617 (0.076) | 0.617 (0.073) |
| **THBS2** | 0.553 (0.071) | 0.579 (0.076) | 0.580 (0.103) | 0.561 (0.097) | 0.581 (0.093) | 0.579 (0.089) |
| **COL1A2** | 0.538 (0.083) | 0.606 (0.062) | 0.616 (0.056) | 0.612 (0.043) | 0.631 (0.028) | 0.630 (0.042) |
| **COL1A1** | 0.531 (0.102) | 0.612 (0.037) | 0.628 (0.051) | 0.619 (0.031) | 0.640 (0.033) | 0.639 (0.042) |
| **COL3A1** | 0.517 (0.072) | 0.590 (0.051) | 0.596 (0.054) | 0.593 (0.035) | 0.612 (0.016) | 0.613 (0.034) |
| **SPARC** | 0.509 (0.069) | 0.592 (0.069) | 0.602 (0.073) | 0.595 (0.077) | 0.616 (0.067) | 0.615 (0.079) |
| **MXRA5** | 0.501 (0.083) | 0.527 (0.093) | 0.521 (0.116) | 0.499 (0.115) | 0.514 (0.104) | 0.508 (0.099) |
| **LUM** | 0.500 (0.073) | 0.595 (0.041) | 0.600 (0.044) | 0.574 (0.047) | 0.600 (0.064) | 0.609 (0.055) |
| **POSTN** | 0.474 (0.110) | 0.484 (0.062) | 0.494 (0.079) | 0.488 (0.051) | 0.506 (0.082) | 0.517 (0.051) |
| **CTSK** | 0.463 (0.051) | 0.509 (0.096) | 0.505 (0.075) | 0.501 (0.099) | 0.522 (0.074) | 0.519 (0.081) |
| **COL5A2** | 0.425 (0.072) | 0.542 (0.032) | 0.544 (0.059) | 0.534 (0.051) | 0.546 (0.050) | 0.547 (0.051) |
| **CTHRC1** | 0.348 (0.190) | 0.438 (0.159) | 0.441 (0.149) | 0.405 (0.189) | 0.441 (0.191) | 0.440 (0.164) |
| **MMP14** | 0.309 (0.094) | 0.417 (0.079) | 0.398 (0.103) | 0.382 (0.080) | 0.391 (0.110) | 0.389 (0.073) |
| **FN1** | 0.282 (0.106) | 0.438 (0.065) | 0.448 (0.078) | 0.452 (0.074) | 0.449 (0.085) | 0.456 (0.064) |
| **ISG15** | 0.241 (0.116) | 0.361 (0.070) | 0.351 (0.090) | 0.352 (0.077) | 0.351 (0.086) | 0.349 (0.068) |
| **COL8A1** | 0.218 (0.032) | 0.321 (0.063) | 0.320 (0.061) | 0.284 (0.037) | 0.314 (0.065) | 0.306 (0.051) |
| **COL10A1** | 0.187 (0.040) | 0.338 (0.045) | 0.334 (0.068) | 0.311 (0.036) | 0.333 (0.069) | 0.328 (0.054) |
| **PDGFRL** | 0.165 (0.093) | 0.231 (0.076) | 0.232 (0.031) | 0.200 (0.070) | 0.224 (0.050) | 0.229 (0.050) |
| **SULF1** | 0.164 (0.032) | 0.303 (0.046) | 0.298 (0.065) | 0.275 (0.032) | 0.300 (0.059) | 0.303 (0.045) |
| **DPYSL3** | 0.155 (0.032) | 0.274 (0.059) | 0.264 (0.072) | 0.232 (0.032) | 0.253 (0.062) | 0.249 (0.049) |
| **IL32** | 0.091 (0.090) | 0.181 (0.099) | 0.176 (0.126) | 0.136 (0.057) | 0.166 (0.120) | 0.168 (0.105) |

**Supplementary Table S1** Semi-supervised learning. The table shows mean and SD values of Pearson's coefficients between the measured and predicted expression levels of 21 breast cancer-related microenvironment marker genes. The 5-fold cross-validation of semi-supervised learning was performed using sections D1 and D3 as training data, section D2 as test data, and sections A–C as unlabeled data.

| Section name | Original image size [pixel] | Spot image size [pixel] | #Measured spots |
|---|---|---|---|
| **Section A** | 11,953 × 12,336 | 80 × 80 | 2,099 |
| **Section B** | 11,845 × 12,011 | 80 × 80 | 2,228 |
| **Section C** | 11,737 × 12,335 | 80 × 80 | 2,479 |
| **Section D1** | 11,501 × 11,849 | 80 × 80 | 2,500 |
| **Section D2** | 11,451 × 12,499 | 80 × 80 | 2,474 |
| **Section D3** | 11,727 × 11,774 | 80 × 80 | 2,361 |

**Supplementary Table S2** Tissue section information. The table shows the original tissue section image size, spot image size, and number of spots measured with Visium platform in sections A–C and D1–D3.
